# Triple antimicrobial combinations with potent synergistic activity against *M. abscessus*

**DOI:** 10.1101/2025.02.13.638181

**Authors:** Yanqin Huang, Lucius Chiaraviglio, Ibidunni Bode-Sojobi, James E. Kirby

## Abstract

Synergy of antimicrobial combinations was tested against contemporary *Mycobacteroides* abscessus isolates using 2-D and 3-D checkerboard assays. Triple combinations of omadacycline-azithromycin-clofazimine, tigecycline-azithromycin-clofazimine, omadacycline-azithromycin-linezolid and omadacycline-azithromycin-contezolid demonstrated synergy (FIC ≤ 0.5) against 33%, 31%, 62%, and 66% of isolates, respectively. Notably, in all triple combinations, macrolide resistant *M. abscessus* subsp. *abscessus* and *M. abscessus* subsp. *bolletii* isolates were fully sensitized to azithromycin at the FIC index, as were isolates with elevated clofazimine MICs.

---

*Mycobacteroides abscessus* complex causes chronic and potentially fatal lung infection especially in patients with underlying structural lung diseases including bronchiectasis, emphysema, and cystic fibrosis [1]. Recommended combinations for initial treatment include at least three active antimicrobials, incorporating a macrolide when active. However, intrinsic resistance to multiple antibacterial classes makes treatment challenging.

The *M. abscessus* complex is divided into three subspecies: *M. abscessus* subsp. *abscessus, M. abscessus* subsp. *bolletii*, and *M. abscessus* subsp. *massiliense. M. abscessus* subsp. *abscessus*, the most common subspecies isolated from human infection, and *M. abscessus* subsp. *bolletii* often express an inducible erythromycin ribosomal methylase (Erm(41)), which confers resistance to macrolides. Moreover, amikacin, an aminoglycoside, has long been a preferred agent in combination regimens. However, kidney damage and permanent hearing loss associated with its long-term use is highly problematic. Therefore, there is an urgent need to optimize treatment regimens.^30^

Several existing antimicrobials that could be used in combinations are noteworthy: **(1)** Omadacycline and tigecycline are third-generation tetracycline class bacterial protein synthesis inhibitors. Several *M. abscessus* murine pneumonia model studies demonstrated efficacy of omadacycline as a single agent and when combined with additional antibiotics [2-5], while *in vitro* synergy with several antibiotics including macrolides was demonstrated against an eleven isolate test set [5]. Two recent large case series described long-term clinical success against *M. abscessus* lung infection in patients treated with omadacycline combined with other agents, as well its safety and tolerability when used in these regimens [6, 7]. **(2)** Linezolid and contezolid are oxazolidinone protein synthesis inhibitors. The latter is currently in phase III clinical trials in the US (NCT05369052, NCT03747497) and is approved for use in China [8]; it also may be associated with lower bone marrow toxicity than linezolid [9, 10]. **(3)** Clofazimine is an inhibitor of mycobacterial DNA replication. There are now several *in vitro* studies also supporting high potency clofazamine against *M. abscessus* [11, 12]. All aforementioned antimicrobials except for tigecycline have an oral dosing option, simplifying long-term therapy.

Several studies have tested antibiotic combinations for *in vitro* evidence of synergy against *M. abscessus* using checkerboard arrays with variable results [5, 13-17]. This variability is likely due in part to the limited numbers of isolates and combinations evaluated in each study. Therefore, here, we examined the combinatorial effects of more recently available and/or less toxic antimicrobials against a large number of *M. abscessus* clinical isolates.

Methodology is described in **Supplementary Materials**. The per isolate MIC, subspecies, and specimen source for isolate sets are listed in **Table S1** and **S2**. MIC_50,_ MIC_90_, and MIC ranges for isolates tested are summarized in **Table S3**. Rates of synergy (FICI ≤ 0.5) are shown in **Table S4**. Antimicrobial concentrations for each antimicrobial alone (MIC) and in combination at the FICI for each isolated tested are listed in **Tables S5-S8**. Median and range of FICI values for combinations are summarized in **Table S9**.

Triple combinations examined showed lower median FICI values compared with constituent pairwise combinations. The pairwise combinations of omadacycline-azithromycin, tigecycline-azithromycin, omadacycline-clofazimine, tigecycline-clofazimine, azithromycin-clofazimine, omadacycline-linezolid, omadacycline-contezolid, azithromycin-linezolid, and azithromycin-contezolid demonstrated synergy in 12% to 38% of isolates (FICI ≤ 0.5), while the triple combination of omadacycline-azithromycin-clofazimine, tigecycline-azithromycin-clofazimine, omadacycline-azithromycin-linezolid and omadacycline-azithromycin-contezolid demonstrated synergy against 33%, 31%, 62%, and 66% of isolates, respectively.

In omadacycline or tigecycline + azithromycin-clofazimine triple combinations, the azithromycin concentration at the FICI was reduced compared with the azithromycin MIC (median concentration 0.25 µg/mL versus 6 µg/mL, or a 24-fold reduction, **Tables S5, S6**). In linezolid or contezolid + omadacycline-azithromycin triple combinations the azithromycin concentration at the FICI was also substantially reduced compared with azithromycin MIC (median concentration 0.125 µg/mL versus 2 µg/mL, or a 16-fold reduction, **Tables S7, S8**). Although many isolates examined were resistant to azithromycin presumptively based on induction of the Erm(41) methylase (i.e. most *M. abscessus* subsp. *abscessus* and *M. abscessus* subsp. *bolletii*), the azithromycin inhibitory concentrations at the FICI in double or triple combinations with omadacycline, tigecycline, and/or clofazimine were ≤ 1 µg/mL, with more pronounced reduction in triple combinations. Therefore, combinations appear to prevent expression or overcome the consequences of inducible macrolide resistance. Induction of Erm(41) methylase has been found to decrease overall bacterial fitness in other contexts based on its perturbation of ribosome function [18]. The observed contribution of azithromycin to synergistic activity in macrolide resistant isolates may therefore also reflect compromised fitness, a hypothesis that would need to be validated through genetic experiments. Taken together, our results suggest that use of macrolides in combination regimens may contribute to antimicrobial activity, even in the presence of resistant macrolide MICs.

In triple combinations, clofazimine inhibitory concentrations at the FICI were also reduced to ≤ 0.25 µg/mL even for isolates with clofazimine MICs as high as 16 µg/mL (**Tables S5, S6**). For context, we note limited clinical reports of clofazimine efficacy for nontuberculous mycobacteria isolates with an MIC ≤ 0.25 µg/mL [19], steady-state concentrations of clofazimine in blood exceeding 0.25 µg/mL [20], and further concentration in macrophages, the presumptive intracellular growth niche.

Notably, the activity of contezolid, an oxazolidinone with potentially lower bone marrow toxicity [9], was comparable to linezolid in all combinations tested. Antagonism was very rare to absent for all combinations examined (**Tables S5-S8**). Of interest, recent murine model studies [3, 4] support therapeutic activity of the omadacycline + clofazimine + linezolid synergistic combination identified in our checkerboard testing, highlighting the utility of checkerboard testing method as a screening tool for identifying combinations worthy of further investigation.

In summary, our data identify triple combinations with compelling activity in checkerboard synergy analysis. Antimicrobial concentrations in triple combinations at the FICI were all ≤ 1 ug/mL for every antimicrobial for all isolates examined including isolates with highly elevated azithromycin and clofazimine MICs. Therefore, several antimicrobial combinations were identified with broad-spectrum *in vitro* activity against *M. abscessus* including those with oral dosing options and which may prove effective against infection.

## Author Notes

We thank Thea Brennan-Krohn, Jessica Pierce, and Kelly Wright for critical reading of the manuscript. J.P. and K.W. are employees of Paratek Pharmaceuticals, Inc.

TECAN and Paratek Pharmaceuticals, Inc., had no role in study design, data collection or interpretation, and decision to publish. Y.H. was a former employee at MicuRx Pharmaceutical Co. (Shanghai, China) and participated in the development of contezolid; she has no financial conflict of interest with the current work.

## Author contributions

Y.H.: conceptualization, methodology, investigation, formal analysis, and manuscript draft and editing. L.C. and I.B-S: investigation. J.K: conceptualization, methodology, formal analysis, and manuscript draft and editing. Funding acquisition was as indicated.

