## Supplemental Materials for "Triple antimicrobial combinations with potent synergistic activity against *M. abscessus*"

**Index:**

**Supplemental Materials and Methods.**

**Supplemental Results and Discussion.**

**Supplemental References.**

**Supplemental Figure S1.**

**Supplemental Tables found in separate Excel spreadsheet**

### Supplementary Methods.

**Antibiotics.** Clofazimine (Acros Organics, Geel, Belgium, Cat# 461760010, lot# A0395020) and omadacycline (Paratek, King of Prussia, PA, lot# CA22-1028) were used for single, pairwise, and triple antimicrobial susceptibility testing of omadacycline, azithromycin, and clofazimine. Omadacycline (TOKU-E, Tokyo, Japan, cat# O048-10mg, lot# 0048-01), linezolid (Acros Organics, Geel, Belgium, cat# 460592500, lot# A0400349), and contezolid (MedChem Express, Monmouth Junction, NJ, cat# HY-19915, lot# 117216,) were used for single, pairwise, and triple antimicrobial susceptibility testing of omadacycline, azithromycin, and linezolid/contezolid. Tigecycline was from Chem-Impex International (Wood Dale, IL, cat# 29737, lot# 001926,) and azithromycin was from Sigma-Aldrich (St. Louis, MO, cat# RHR1088-1G, lot# LRAB3758).

**Bacterial isolates**. Clinical isolates were obtained from patient samples at Beth Israel Deaconess Medical Center (Boston, MA, USA) from 2016-2023 under an Institutional Review Board-approved protocol; BEI Resources (Manassas, VA); and the American Type Culture Collection (Manassas, VA). *M. peregrinum* ATCC 700686 and *S. aureus* ATCC 29213 were used as quality control strains based on Clinical Laboratory and Standards Institute (CLSI) guidance [1]. *M. abscessus* were subspeciated using the method of Akwani et al. [2]. All isolates and strains were stored at -80°C and minimally passaged until use in experiments.

**Minimal inhibitory concentration testing.** Broth microdilution antimicrobial susceptibility testing was performed per Clinical Laboratory and Standards Institute (CLSI) protocol. Serial two-fold dilutions of freshly prepared drugs were dispensed using the HP D300 digital dispensing system (HP, Inc., Palo Alto, CA) into sterile, round-bottom, polystyrene 96-well plates (Evergreen Scientific, Los Angeles, CA) to achieve final desired antimicrobial concentrations [3].

Inocula were prepared by suspending colonies in normal saline (0.9% NaCl; Thermo Fisher Scientific, Waltham, MA) and adjusting to 0.5 McFarland standard using a DensiChek Plus handheld colorimeter (bioMérieux, Durham, NC). The resulting suspensions was diluted 1:200 in freshly prepared cation-adjusted Mueller-Hinton broth (BD Diagnostics, Franklin Lakes, NJ). 100 µL of each suspension was added to each well using a Welljet reagent dispenser (Integra Biosciences, Hudson, NH) to achieve a final bacterial inoculum of approximately $5\times10^5$ CFU/mL. Following incubation at 30°C in ambient air, minimal inhibitory concentration (MIC) results were recorded from day 3 to day 5, except for azithromycin, which was recorded on day 14 [1].

Quality control was performed per CLSI guidelines [4] and fell within acceptable ranges as available (i.e., for linezolid, omadacycline, tigecycline) for all experiments. Paratek and TOKU-E omadacycline quality control minimal inhibitor concentration (MIC) values were within one doubling dilution of one another in all experiments [1].

Analysis was performed using two overlapping sets of clinical isolates and antibiotics with the distinction that omadacycline was obtained either from the manufacturer (Paratek) of the FDA-approved drug in the first set of synergy experiments and from a commercial supplier ( TOKU-E) in the second set of synergy experiments as delineated in Supplementary Tables. The isolates tested in the second set of synergy experiments (n=29) were selected randomly from the isolates (n=51) tested in the first set of synergy experiments. The per isolate MIC, subspecies, and specimen source in these isolate sets are listed in **Table S1** and **Table S2**, respectively.

**Synergy assays.** To set up synergy testing, drugs combinations were dispensed using the HP D300 digital dispensing system (HP, Inc., Palo Alto, CA) as previously described [3]. To limit the number of potential combinatorial permutations, the two or three drugs in each combination were added together in a doubling dilution series starting at or above the individually determined MICs for each drug with an additional seven to nine doubling dilutions below this, as described previously [5]. Testing of individual drug MICs was performed concurrently. Using this method, the concentrations of antimicrobials in the combinations examined lie on the diagonal of a square or cubic checkerboard synergy grid. Incubation conditions were the same as for single drug testing, with all combinations containing azithromycin read on day 14.

To test for drug interaction, the fractional inhibitory concentration index (FICI) was determined by summing the individual fractional inhibitory concentrations (FICs) for the two or three antimicrobials in each inhibited well, where the FIC equals the inhibitory concentration of the individual drug in the combination divided by the MIC of the drug when tested alone. The scored FICI was the lowest summation for which complete visual growth inhibition was observed. FICI values of $\leq 0.5$, $0.5 < \text{FICI} \leq 4.0$, and $\text{FICI} > 4.0$ were scored as indicating synergy, indifference, or antagonism, respectively [6].

**Supplemental Results and Discussion.**

Interestingly, three classes of protein synthesis inhibitors exhibited synergy with one another: specifically, tetracyclines, which bind to the A-site decoding center of the 30S ribosomal subunit and oxazolidinones and azithromycin, which binds to the 50S subunit (**Fig. S1A**). Superimposition of cryo-EM structures of the two oxazolidinones examined and azithromycin bound to the *E. coli* ribosome show that these agents respectively bind in adjacent regions of the

23S rRNA separated by approximately 3Å (**Fig. S1B**). It is unclear whether their adjoining binding sites promote cooperative binding, allowing occupancy of only one of the two classes of antibiotics at their binding sites, or neither. Of note, oxazolidinones and azithromycin functionally block the peptidyl transferase center and nascent peptide exist tunnel, respectively, potentially allowing interference with functionally distinct aspects of protein synthesis. The synergy observed between azithromycin and oxazolidinones may result from direct (e.g., cooperative binding, functional synergy) or indirect mechanisms (e.g., effects on *erm(41)* gene expression), both of which could explain our observations.

**Supplemental References.**


1. Anonymous. 2023. CLSI. Performance Standards for Susceptibility Testing of Mycobacteria, Nocardia spp., and Other Aerobic Actinomycetes. 2nd edition. CLSI supplement M24S. Clinical and Laboratory Standards Institute; 2023.
2. Akwani WC, van Vliet AHM, Joel JO, Andres S, Diricks M, Maurer FP, Chambers MA, Hingley-Wilson SM. 2022. The Use of Comparative Genomic Analysis for the Development of Subspecies-Specific PCR Assays for Mycobacterium abscessus. Front Cell Infect Microbiol 12:816615.
3. Brennan-Krohn T, Truelson KA, Smith KP, Kirby JE. 2017. Screening for synergistic activity of antimicrobial combinations against carbapenem-resistant Enterobacteriaceae using inkjet printer-based technology. J Antimicrob Chemother 72:2775-2781.
4. Anonymous. CLSI. Performance Standards for Susceptibility Testing of Mycobacteria, Nocardia spp., and other Aerobic Actinomycetes. 1st ed. CLSI supplement M62. Wayne, PA: Clinical and Laboratory Standards Institute; 2018.
5. Berenbaum MC, Yu VL, Felegie TP. 1983. Synergy with double and triple antibiotic combinations compared. J Antimicrob Chemother 12:555-63.
6. Odds FC. 2003. Synergy, antagonism, and what the chequerboard puts between them. Journal of Antimicrobial Chemotherapy 52:1.
7. Pellegrino J, Lee DJ, Fraser JS, Seiple IB. 2022. E. coli 50S ribosome bound to tiamulin and azithromycin. PDB 8E42.https://doi.org/10.2210/pdb8E42/pdb.
8. Paternoga H, Crowe-McAuliffe C, Bock LV, Koller TO, Morici M, Beckert B, Myasnikov AG, Grubmüller H, Nováček J, Wilson DN. 2023. Structural conservation of antibiotic interaction with ribosomes. Nature Structural & Molecular Biology 30:1380-1392.
9. Tsai K, Stojković V, Lee DJ, Young ID, Szal T, Klepacki D, Vázquez-Laslop N, Mankin AS, Fraser JS, Fujimori DG. 2022. Structural basis for context-specific inhibition of translation by oxazolidinone antibiotics. Nature Structural & Molecular Biology 29:162-171.
10. Wright A, Deane-Alder K, Marschall E, Bamert R, Venugopal H, Lithgow T, Lupton DW, Belousoff MJ. 2020. Characterization of the Core Ribosomal Binding Region for the Oxazolidone Family of Antibiotics Using Cryo-EM. ACS Pharmacol Transl Sci 3:425-432.

**Supplemental Figure S1. Binding sites of ribosome-targeting antibiotics tested in synergy experiments. (A)** Linezolid (orange) and azithromycin (yellow) bind in the peptidyl transferase center and peptide exit tunnel, respectively, in the 50S ribosomal subunit of the bacterial 70S ribosome. Omadacycline binds within the A-site decoding center in the 30S ribosomal submit. Image is a superimposition of PDB 8E42[7], 8CA7[8], and 7S1G[9]. **(B)** Oxazolidinones, linezolid (purple) and contezolid (black), overlap and bind adjacent to azithromycin (yellow), separated by approximately 3Å. 23S rRNA nucleotide A-2058 (cyan) becomes dimethylated by erm(41), blocking binding of macrolides through steric interference. Proximity of A-2508 to the azithromycin desosamine sugar can be appreciated. The panel is a superimposition of PDB 8E42, 7S1G and 6WQN[10]. PDB alignment and image generation were performed using Pymol 2.5.2 (Schrodinger, LLC).

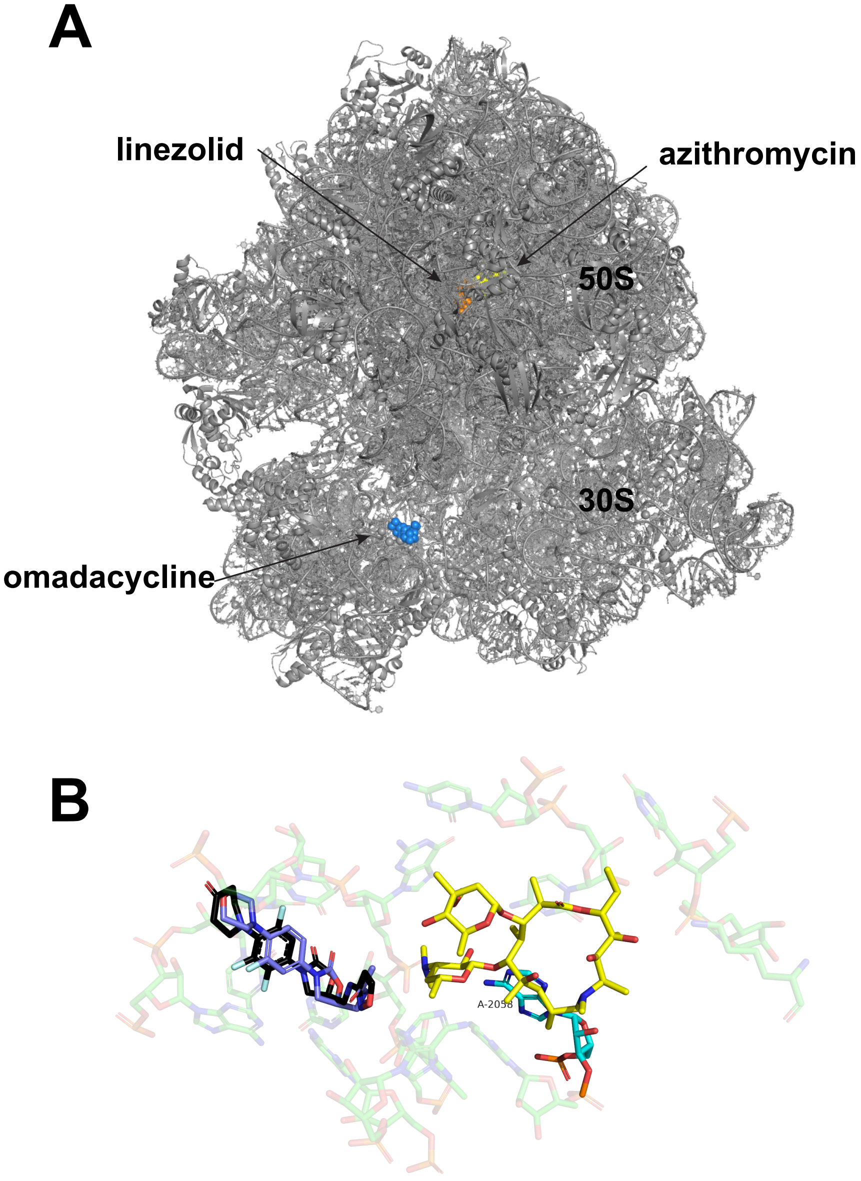
